# Annotation-Free Deep Learning for Predicting Gene Mutations from Whole Slide Images of Acute Myeloid Leukemia

**DOI:** 10.1101/2023.11.13.563550

**Authors:** Bo-Han Wei, Xavier Cheng-Hong Tsai, Kuo-Jui Sun, Min-Yen Lo, Sheng-Yu Hung, Wen-Chien Chou, Hwei-Fang Tien, Hsin-An Hou, Chien-Yu Chen

## Abstract

The rapid development of deep learning in recent years has revolutionized the field of medical image processing, including the applications of using high-resolution whole slide images (WSIs) in acute myeloid leukemia (AML) diagnosis. Although the potential of characterizing gene mutations directly from WSIs has been demonstrated in some cancers, it still faces challenges due to image resolutions and manual annotations. To address this, we propose a deep learning model based on multiple instance learning (MIL) with ensemble learning to predict gene mutations from AML annotation-free WSIs. Our deep learning model offers a promising solution for gene mutation prediction on *NPM1* mutations and *FLT3* -ITD without the need for patch-level or cell-level manual annotations, reducing the manpower and time costs associated with traditional supervised learning approaches. The dataset of 572 WSIs from AML patients that we used to train our MIL models is currently the largest independent database with both WSI and genetic mutation information. By leveraging upsampling and ensemble learning techniques, our final model achieved an AUC of 0.90 for predicting *NPM1* mutations and 0.81 for *FLT3* -ITD. This confirms the feasibility of directly obtaining gene mutation data through WSIs without the need for expert annotation and training involvement. Our study also compared the proportional representation of cell types before and after applying the MIL model, finding that blasts are consistently important indicators for gene mutation predictions, with their proportion increasing in mutated WSIs and decreasing in non-mutated WSIs after MIL application. These enhancements, leading to more precise predictions, have brought AML WSI analysis one step closer to being utilized in clinical practice.

## 1. Introduction

Acute myeloid leukemia (AML), as an aggressive hematologic malignancy, exhibits significant biological and clinical heterogeneity, characterized by uncontrolled proliferation and impaired differentiation of hematopoietic precursors [1]. For optimizing treatment efficacy and minimizing treatment-related complications, precise risk stratification stands as the cornerstone. Currently, a range of cytogenetic changes and gene mutations (expressions) have been integrated into the risk stratification, shaping the treatment landscape [2,3]. Nucleophosmin 1 (*NPM1*) and FMS-like tyrosine kinase-3 internal tandem duplication (*FLT3* -ITD) are the most prevalent recurrent gene mutations in patients with AML [4,5]. Considerable efforts have been devoted to developing targeted therapies against these mutations [6,7], highlighting their crucial significance in clinical practice. However, performing molecular testing for these mutations presents notable challenges; it is labor-intensive, expensive, and often difficult to provide timely results.

Whole slide images (WSIs) employ digital imaging technology to transform pathological specimens into high-resolution digital images detailing cellular and histological structures [8]. While deep learning has shown promise in tasks such as binary morphological classification and histological grading using WSIs [9,10], challenges persist in analyzing bone marrow aspirates due to their complex cytological nature. Aspirates typically feature small, cluttered regions with various cell types and non-cellular debris. Identifying regions of interest (ROIs) and distinguishing individual cells or objects from the background require multi-step preprocessing, including segmentation and denoising [11,12]. Despite advances in deep learning for object detection, manual annotation of segmented cells by experts remains labor-intensive and time-consuming [13]. Developing a more efficient and accurate method for analyzing bone marrow aspirate WSIs is essential.

Contemporary histopathology research often follows a two-stage workflow, focusing on patch-level and slide-level training [14]. Initially, a Convolutional Neural Network (CNN) is trained on patches extracted from WSIs with patch-level annotations, learning complex patterns. In the second stage, features learned at the patch level are utilized to train a slide-level model, necessary for diagnosis from WSIs, capitalizing on insights from patch-level analysis. Widely used for cancer identification [15–17], classification [18,19], and metastasis detection [15], these approaches require substantial manual annotations. Multiple Instance Learning (MIL) has been employed to directly use slide-level labels [20,21], reducing the annotation burden by classifying slides based on the highest-scoring patch.

We hypothesized that deep learning could predict gene mutations based on cellular morphology. Here, we present an end-to-end artificial intelligence framework for bone marrow cytology, uniquely trained using WSIs with slide-level annotations. By leveraging annotation-free WSIs for gene mutation predictions, we demonstrate the capability of deep learning in predicting gene mutations. Our results highlight that models trained at the cell level outperform those trained at the patch level. Additionally, we illustrate how techniques, such as upsampling and ensemble learning, can enhance the predictive performance of the model, especially in scenarios with limited training data.

## 2. Results

### 2.1. Automatic Selection of ROI Patches

In digital pathology, glass slides of bone marrow aspirate smears were scanned using a digital slide scanner to generate high-resolution WSIs for hematopathologist analysis (Fig. 1a). To initiate this process, a dataset was sampled from 572 bone marrow aspirate WSIs obtained at the National Taiwan University Hospital. To address the issue of detecting ROI patches, we developed a pipeline to select ROI patches in stages, automatically identifying areas within the bone marrow aspirate WSIs that are suitable for cytological analysis.

**Figure 1.**
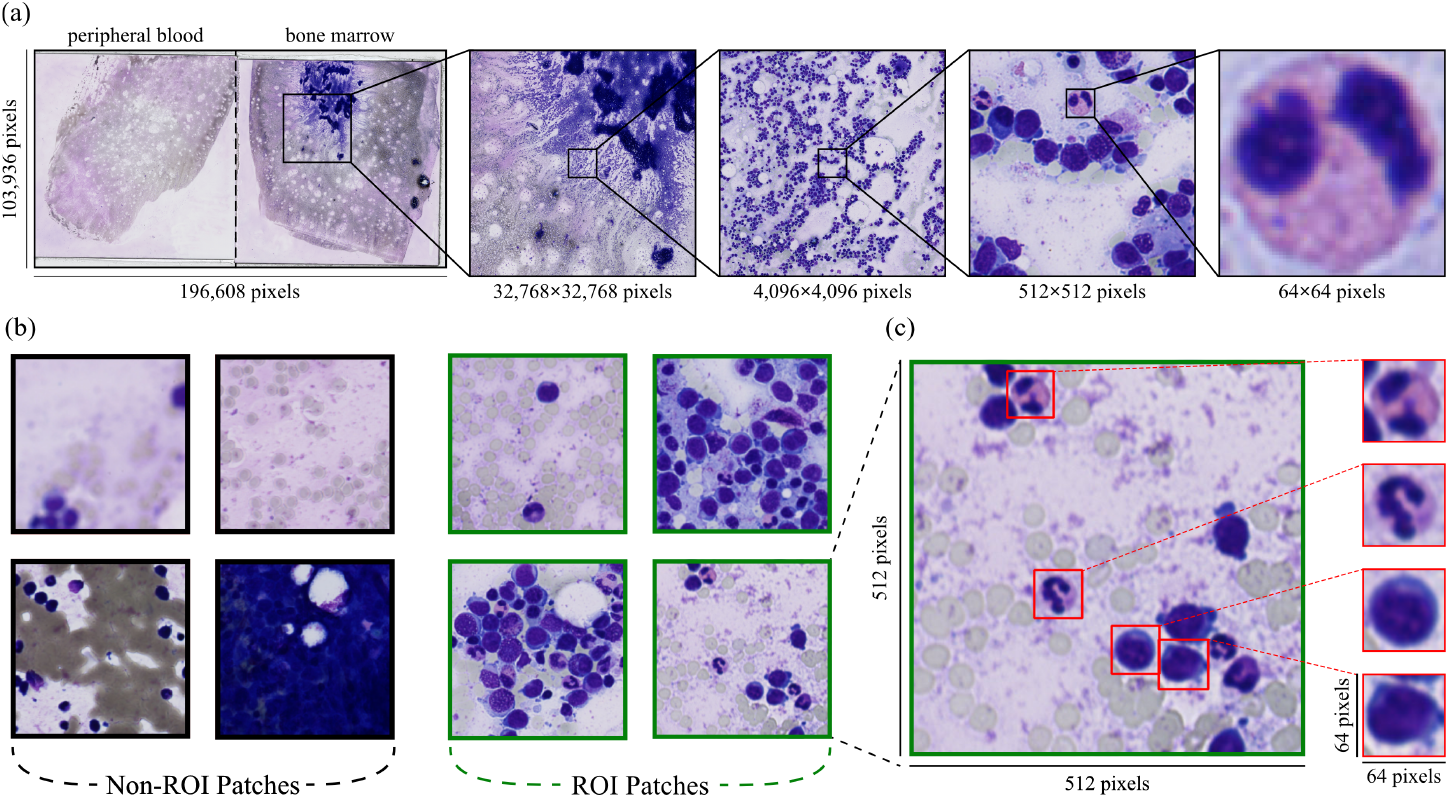
Different scales of images. **a**. The WSI has an original resolution of approximately 200,000 × 100,000 pixels. **b**. ROI patches cover areas with single or multiple leukocytes that are evenly distributed, clear, and not blurry. Non-ROI patches contain only red blood cells but no leukocytes, regions with cell overlaps making identification challenging, a considerable mix of heterogeneous tissues, areas devoid of blood cells within the designated range, or blurry patches. **c**. The resolution of the patches is 512×512 pixels. The size of filtered cells is within the range of 51×51 pixels to 80×80 pixels and then resized to a uniform size of 64×64 pixels.

A single WSI might contain only a few areas suitable for cytology. These areas are sparsely distributed, with minimal cell overlap and staining artifacts, and exhibit subtle and intricate cytological features required for cell classification. To acquire such areas efficiently, the PyHIST tool [22] was first used with the default graph method to remove the blank background regions, retaining only the stained portions. Subsequently, we employed a fine-tuned DenseNet121 architecture, utilizing pre-trained weights from previous research [23] to classify individual patches as ROI patches or non-ROI patches (Fig. 1b).

We observed that the classification results aligned with expectation, i.e., a real-world scenario where usually only 10-20% of a WSI might be the ROI regions for cytology. The results obtained after applying DenseNet121 reveal that for most WSIs, the ROI patches were reduced to around 10-25% of their original counts after undergoing this selection process, indicating a substantial reduction in patch counts (Fig. S1). This screening process significantly reduced the time required for subsequent cell detection processes and exhibited outstanding filtering effectiveness in removing areas with excessive cell overlap, excessive tissue artifacts, or patches devoid of blood cell presence within the designated range. These problematic patches were effectively eliminated, while patches with a minimal number of cells were not erroneously dropped (Fig. 1b).

### 2.2. Leukocyte Detection

After the ROI selection process, we employed a YOLOv4 model to automatically detect and classify cells and non-cellular objects within the selected ROI patches. The training weights in the same study of selecting ROI patches [23] were used to fine-tune this model, with ROI patches identified by the ROI detection model as input. The YOLOv4 model aimed to automatically detect and classify all bone marrow cellular and non-cellular objects. We further set a confidence threshold of 0.5. Cells with confidence scores below this threshold were not captured, ensuring high-quality cell selection for subsequent MIL model training. In addition to managing confidence levels, we factored in cell size during the process. Cells were chosen if their sizes were within the range of 51× 51 pixels to 80× 80 pixels, where the patch size is 512 ×512. The chosen cells were then uniformly resized to 64 ×64 pixels, ensuring a consistent input data size for convenient execution of subsequent augmentation steps (Fig. 1c).

The number of cells left per WSI varied between 100 to 100,000, with the majority of cell counts being below 20,000 and an average of 11,273 (Fig. S2). This outcome effectively reduced the input quantity for the MIL model while maintaining the best quality of all input cells, accelerating training time without compromising accuracy. Therefore, we used leukocytes, including basophil, blast, eosinophil, lymphocyte, metamyelocyte, monocyte, myelocyte, neutrophil, and promyelocyte as inputs for MIL training, denoted as ‘all cells’. Among these cells, MIL randomly selected 2,000 cells as representative cells of a WSI, i.e., a bag. During the selection period, we applied the upsampling technique to the mutated WSIs to balance the data. By separating each mutated WSI into multiple bags, we can generate numerous mutated bags that constitute 2000 cells. For mutated and standard bags with fewer than 2,000 cells, we applied image augmentation to the cells within these bags, which ensured the minimum number of cells was 2,000 in each bag and enhanced the model’s robustness.

### 2.3. Cell-Level Multiple Instance Learning

The total dataset of 572 WSIs was split into training, validation, and test sets at slide level, with a ratio of around 7:1: 2 (400:56:116 WSIs). We used the DenseNet121 as the base embedding model, initialized with pre-trained weights from ImageNet. The MIL model underwent training for 100 epochs with a learning rate set to 0.0001. The loss minimization is attained through stochastic gradient descent (SGD) utilizing the Adam optimizer. The batch size is determined by the number of cells within each bag, restricted to a maximum of 2000 instances (cells) per batch. Since the training data still exhibited minor class imbalance after upsampling we set weights (*w*0, *w*1) to (0.49, 0.51) for *NPM1* mutations and (0.63, 0.37) for *FLT3* -ITD to address this issue. To emulate a real-world scenario, each epoch was evaluated using an imbalance validation set of 56 WSIs, specifically, more standard than mutated WSIs. Early stopping was implemented to prevent overfitting. The separate testing set included 96 standard and 20 *NPM1* mutations WSIs, along with 95 standard and 21 *FLT3* -ITD WSIs.

In addition to established MIL training, we leveraged ensemble learning by employing different values of *K*( from *K* = 1 to *K* = 30) in MIL to diversify the models’ perspectives on the data. Each *K* represents the number of positive instances considered within a bag during the MIL training process, leading to the creation of models with varying focuses on distinct data subsets. Based on Eq. 2 and Eq. 3, we calculated the respective weights 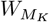 of all MIL models and then estimated the final prediction. The MIL integrated with ensemble learning and upsampling techniques demonstrated its efficacy and surpassed the traditional MIL training at both the patch level and cell level. For instance, the cell-level MIL we proposed achieved the highest classification performance for *NPM1* mutations with an AUC of 0.90, and this method also revealed an AUC of 0.81 for *FLT3* -ITD (Fig. 2). This achievement is better than that of Kockwelp et al. [13]. We consider this owing to the proposed cell inference and ensemble strategies. The performance on *NPM1* is comparable to the strategy based on cell-level annotation [24].

**Figure 2.**
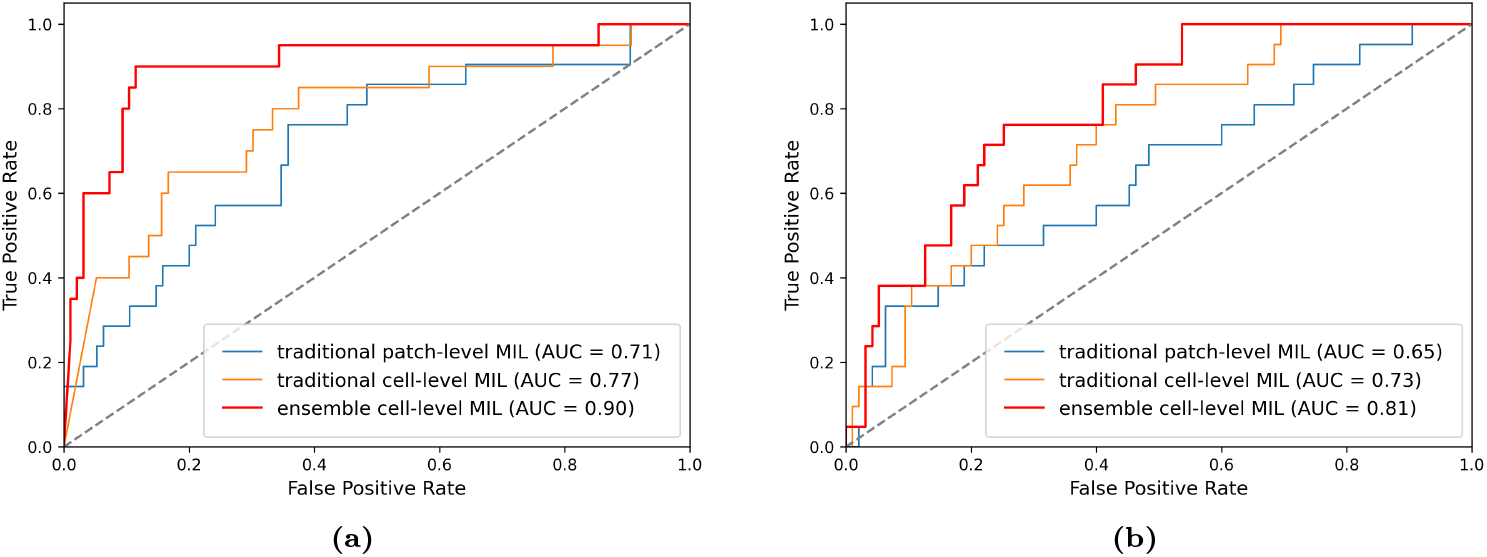
ROC curves for MIL model trained using different strategies. ’Traditional’ refers to MIL without upsampling and ensemble learning, while ‘Proposed’ refers to MIL with upsampling and ensemble learning. **a**. Models trained with upsampling *NPM1* mutations data. **b**. Models trained with upsampling *FLT3* -ITD data.

Moreover, our ensemble cell-level MIL, when using sensitivity as the standard for the reliability analysis of clinical diagnosis, exhibited better performance than the traditional MIL methods. As the sensitivity was set to 0.75, our approach reduced the false positive rate from 0.36 to 0.09 for *NPM1* mutations. For *FLT3* -ITD, there was also a notable reduction from 0.60 to 0.25. This outcome once again confirms that ensemble learning and upsampling can enhance the effectiveness of the MIL model.

To clarify the importance of cell features for prediction in the learning process of MIL, we compared the proportional representation of nine types of cells. Before applying the MIL model, all cell types in each WSI would be divided by the total number of cells of all types in that WSI to obtain the proportion of each type of cell. After prediction, we listed the top 100 images of cells with the highest probability of being predicted by the model as the highest correlated with mutations in each bag. Then, we calculated the proportion of all representative cells predicted by MIL in the original WSI based on their cell types (Fig. 3). This can reveal which cells are more meaningful for MIL predictions of gene mutations, even if we were only using slide-level annotations.

**Figure 3.**
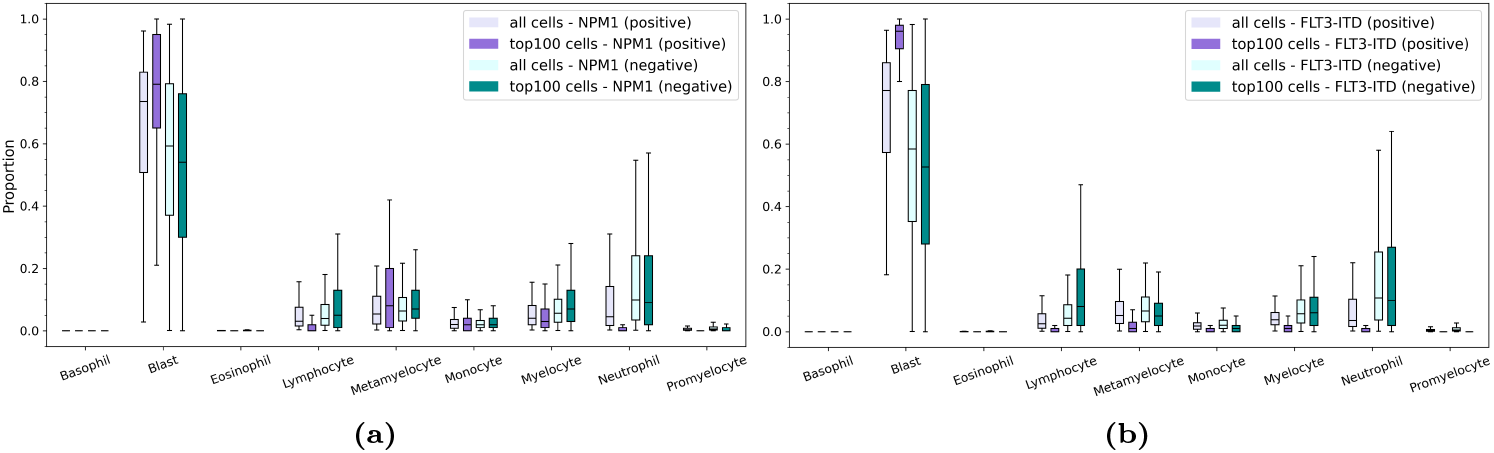
Comparision of cell types. All WSIs were categorized based on their slide-level labeling. ‘Positive’ refers to WSIs with the gene mutation, while ‘negative’ refers to WSIs without the gene mutation, i.e., the standard WSIs. **a**. boxplot of cell types with *NPM1* mutations data. **b**. boxplot of cell types with *FLT3* -ITD data.

It can be observed that the trend for blasts remains consistent regardless of whether it is the *NPM1* or *FLT3* -ITD model. In bags with either genetic mutations, after applying MIL, there’s a significant increase in the proportion of blasts among the top 100 representative cells. On the other hand, in bags without genetic mutations, after applying MIL, there’s a noticeable decrease in the proportion of blasts among the top 100 representative cells. This trend indicates that the presence of blasts is an important indicator for determining whether there are mutations in deep learning. Additionally, in bags with genetic mutations, the proportions of other cell types decrease, indicating that the majority of features used in machine learning to determine the presence of mutations are in the blast category. Therefore, in non-mutated bags, because these features cannot be found in blast cells, the proportions of other cell types increase while the proportion of blast significantly decreases.

## 3. Methods

### 3.1. Dataset

Between 1994 and 2015, 572 patients who were diagnosed with *de novo* AML at the NTUH were enrolled in this study. Bone marrow smears and peripheral blood smears were scanned as WSIs after using a modified Romanowsky stain (Fig. 1a). The gene mutation status was determined using TruSight myeloid panel on the HiSeq platform (Illumina, San Diego, CA) [25], which were annotated as 1 (indicating the presence of pathogenic or likely pathogenic mutations) or 0 (indicating the absence of pathogenic or likely pathogenic mutations). In our cohort, a total of 34 genes were frequently found to be mutated, and in this study, we listed the mutation frequencies of more than 10% and selected the top two highest frequent mutations as the targets (Table 1). This retrospective study was approved by the NTUH Research Ethics Committee, and written informed consent was obtained from all participants in accordance with the Declaration of Helsinki (Approval number: 201802021RINC).

**Table 1.**
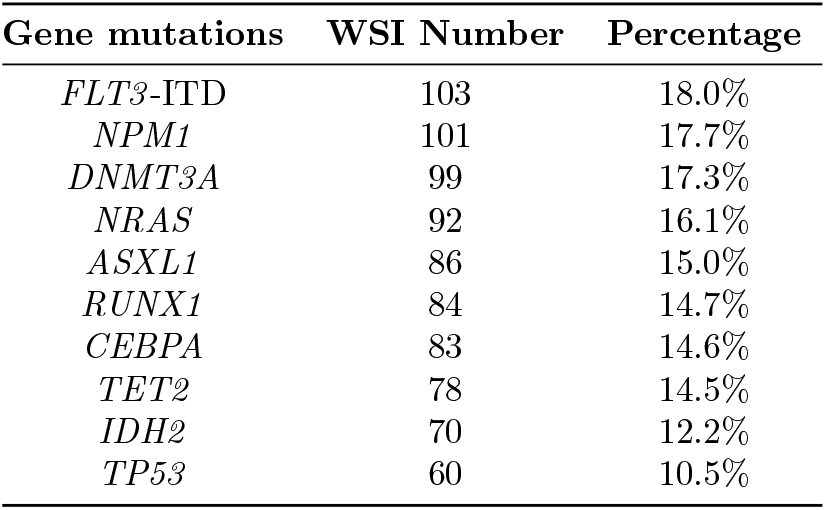
Number of WSIs with gene mutations in our cohort.

| Gene mutations | WSI Number | Percentage |
| --- | --- | --- |
| *FLT3*-ITD | 103 | 18.0% |
| *NPM1* | 101 | 17.7% |
| *DNMT3A* | 99 | 17.3% |
| *NRAS* | 92 | 16.1% |
| *ASXL1* | 86 | 15.0% |
| *RUNX1* | 84 | 14.7% |
| *CEBPA* | 83 | 14.6% |
| *TET2* | 78 | 14.5% |
| *IDH2* | 70 | 12.2% |
| *TP53* | 60 | 10.5% |

### 3.2. Cell Image Generation

The bone marrow smears of WSIs underwent a three-step filtering process to identify the cells for subsequent model training. First, we used the PyHIST tool to generate patches [22]. This tool was applied to filter out background regions and non-smear areas in the WSIs (Fig. 4a). The patches (512 ×512 pixels) were extracted from the highest resolution (40X magnification), and the generation method was based on the graph method, with a content threshold of 0.05. The parameters ‘tilecross-downsample’ and ‘mask-downsample’ adopted the default values, and the parameter ‘output-downsample’ was set to 1 in order to obtain patches at the original resolution. Secondly, the patches generated from the PyHIST tool were further classified into ROI and non-ROI patches. We utilized an ROI detection model from a previous study [23], which was based on DenseNet121 [26] and was pre-trained and fine-tuned on labeled patches (ROI/non-ROI). In this step (Fig. 4b), patches containing densely packed leukocytes or areas without any leukocytes were removed, significantly reducing the number of input data for the subsequent step: cell detection modeling.

**Figure 4.**
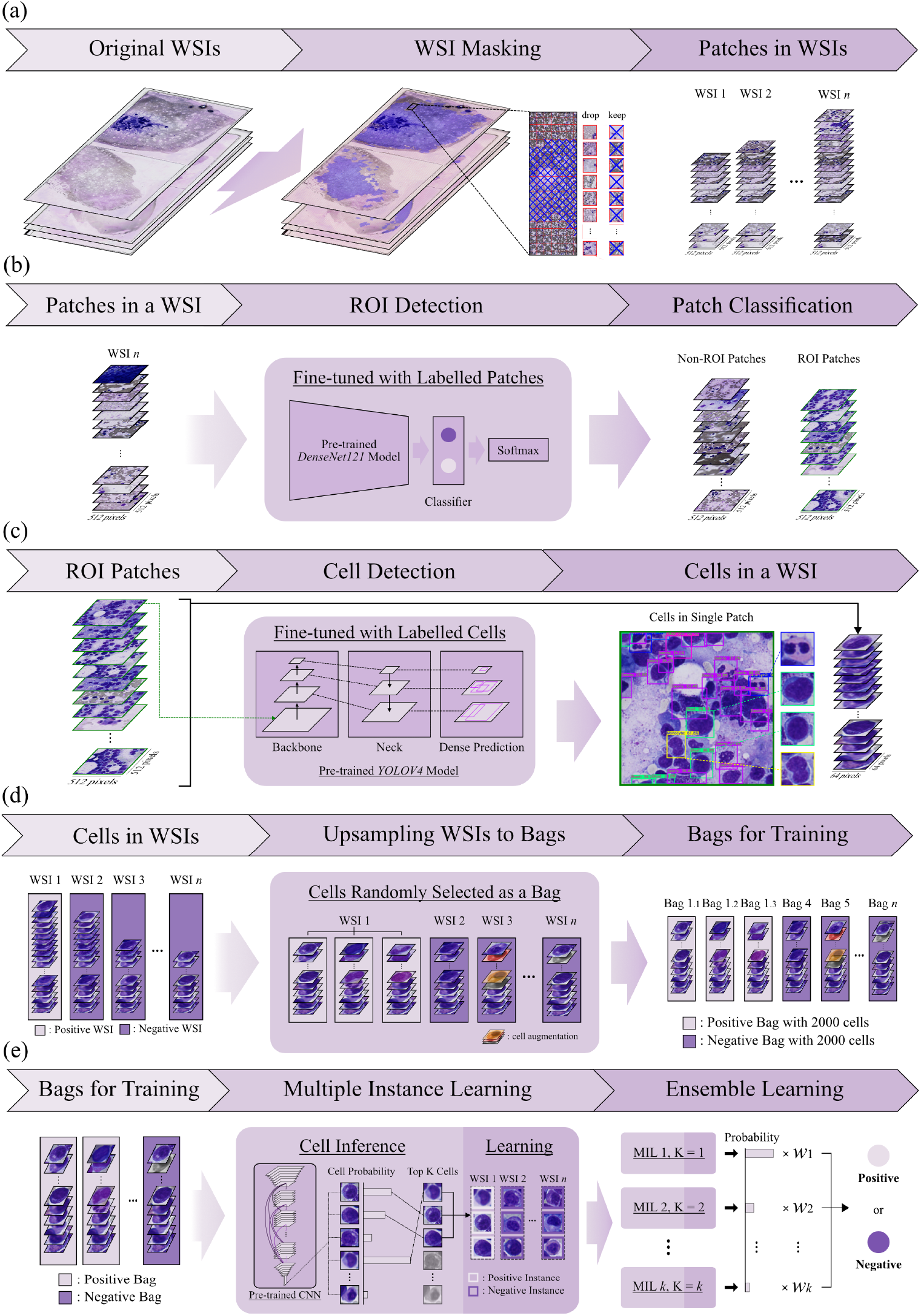
Overview of the proposed method. **a**. Generation of patch images. **b**. Selection of ROI patches. **c**. Leukocyte (cell) detection. **d**. Bag upsampling and cell augmentation processes. **e**. MIL and ensemble learning.

After ROI selection, the ROI patches in each WSI would be used to get the cells in the third step. We applied the cell detection model from the same research of ROI detection [23]. This model was based on YOLOv4 [27] and trained to predict the bounding boxes of leukocyte objects. In this step, all leukocytes in each ROI patch in bone marrow aspirates were detected (Fig. 4c).

### 3.3. Multiple Instance Learning

Fully supervised approaches for histopathology image analysis require detailed manual annotations, which are time-consuming and intrinsically ambiguous, even for well-trained experts. Standard unsupervised approaches usually fail due to their complicated patterns.

MIL is a weakly supervised learning approach used in tasks where the training data consists of labeled groups of instances, known as bags, rather than individually labeled instances. It works well for the current study because it takes advantage of both supervised and unsupervised approaches. The main idea of MIL is to learn local patterns using global annotations. In MIL, each bag contains multiple instances, but only the bag is labeled with a class label, and the instances within the bag are unlabeled. In previous research, they transformed binary classification tasks into MIL problems by dividing WSIs into multiple instances. Here, the instances could be patches [20] or could be cells as proposed in this study. Positive WSIs contain at least one positive instance, while negative WSIs are without any positive instances. With the above two criteria, we expect positive WSIs to contain only a few key instances, while negative WSIs can have all the patches or cells as instances.

In this study, the term “bags” refers to WSIs, while “instances” denotes distinguishable cells. We categorize WSIs with and without mutations as mutated and standard bags, respectively. Each WSI can be seen as a bag containing multiple cells (Fig. 4d). To predict a bag as mutated, at least one cell should be classified as mutated. On the other hand, all cells should be classified as standard when predicting a bag as standard. In a bag, all cells will be classified, and they are ranked based on their probability of being classified as mutated. For mutated bags, the probability of the highest-ranked cell should be higher than a threshold, while for standard bags, it should be lower than that threshold. The MIL task involves learning a cell-level representation that effectively discriminates distinguishing cells within mutated WSIs from all other cells. Based on the cell detection process described above, bags *B* = {*B*_*i*_ : *i* = 1, 2,. .., *n*} are generated, where 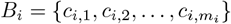 is a bag containing *m*_*i*_ cells from WSI *s*_*i*_.

During training, the embedding models AlexNet, DenseNet, EfficientNet, ResNet34, ResNet101, ResNet152, ResNext, and VGG16 were employed and compared. These models are initialized with pre-trained weights from ImageNet. The model is represented as a function *f*_*θ*_, where the current parameters θ map the input cells *c*_*i,j*_ to probabilities of “standard” and “mutated” classes. For a bag *B*_*i*_, a vector list 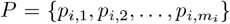 is obtained, with each vector corresponding to the probability of the “mutated” class for each cell *c*_*i,j*_ (*j* = 1, 2, …, *m*_*i*_, where *m*_*i*_ is the number of cells in *B*_*i*_) in WSI *s*_*i*_. Once we have obtained the index *k*_*i*_ of the highest-ranked “mutated” cell for each WSI, denoted as *k*_*i*_ = *argmax*_*j*_ (*p*_*i,j*_),*j* = 1, 2, …, *m*_*i*_, we proceed to label this specific cell as “mutated.” This represents the strictest version of MIL, but the standard MIL assumption can be relaxed by introducing a hyperparameter *K*, assigning the top *K* cells as mutated in mutated WSIs. For *K* = 1, the highest-ranked cell in bag *B*_*i*_ is 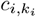.Then, the network’s output 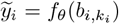 is compared to the label of WSI *y*_*i*_ of WSI *s*_*i*_ using the cross-entropy loss function *l* as in Eq. 1:

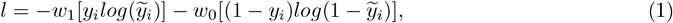

where *w*_0_ and *w*_1_ are the weights for standard and mutated classes, the sum of these two weights must equal 1. Similarly, if *K>* 1, all the top *K* cells from a WSI share the same label *y*_*i*_, and the loss can be computed with Eq. 1 for each one of the *K* cells.

### 3.4. Data Upsampling

The class imbalance in the WSI data was indeed substantial, with a limited number of samples exhibiting gene mutations compared to standard samples, which could significantly impact training outcomes. To address this imbalance, we introduced class weights, denoted as *w*_0_ and *w*_1_ for the standard and mutated classes, respectively. These weights allowed us to assign greater importance to the underrepresented mutated examples.

Furthermore, prior to MIL training, we faced the challenge of working with WSIs, each containing tens of thousands of cells. To expedite the training process, we randomly selected a fixed number (2,000) of cells to represent each WSI, reducing the input quantity. Since the number of mutated WSIs was considerably smaller compared to the negative class, we adopted a strategy of dividing mutated WSIs into multiple bags, each containing the same fixed number of cells. These bags were considered independent training data, effectively increasing the training data for the mutated class and resulting in a more balanced training dataset. For all bags with fewer than 2,000 cells, we applied image augmentation to extend the cell number within these bags. The augmentation included flipping, translation, rotation, and color adjustments such as random contrast (scaling by 0.5–1.5), brightness (scaling by 0.65–1.35), hue (shifting by -32–32), and value (shifting by -32–32).

### 3.5. Ensemble Learning

Ensemble methods represent a potent approach that encompasses the training and amalgamation of multiple models to tackle intricate problems. At its heart, ensemble methods rely on the fundamental concept that a gathering of individual “weak learners” can synergistically give rise to a “strong learner.” In this strategy, each model casts its vote or contributes valuable insights, with the ensemble method harmonizing these inputs to formulate a definitive prediction. The overarching objective of ensembles is to mitigate bias and variance in predictions. By harnessing the combined capabilities of multiple models instead of relying solely on a single one, these methods enhance accuracy and robustness significantly.

In this study, we utilized an ensemble method based on a loss function approach. This technique aimed to optimize the model combination of each base model *M*_*K*_(*K* = 1, 3, 5, 10, 20, where *K* is the top *K* cells) by employing a loss function 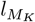 as in Eq. 1 to quantify the difference between predicted and actual values. Each base model within the ensemble was assigned a weight 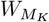 determined by its performance as in Eq. 2:

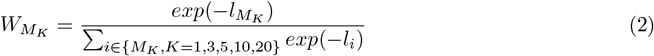

The ensemble model then combined the probabilities 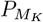 of these models according to their respective weights, as illustrated in Eq. 3:

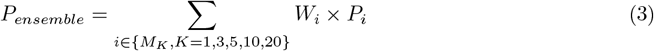

This approach ultimately led to improved prediction accuracy.

## 4. Conclusion

In our research, we extensively explored Multiple Instance Learning (MIL) with upsampling and ensemble techniques to address the challenges in diagnosing acute myeloid leukemia (AML). Our strategy achieved impressive AUC values for *NPM1* mutations, reaching 0.90, and for *FLT3* -ITD, which achieved 0.81. Furthermore, the proposed method aims at maintaining the same sensitivity while reducing false positive occurrences. In *NPM1* mutation prediction, we diminished the false positive rate from 0.36 to 0.09, while for *FLT3* -ITD, the false positive rate dwindled from 0.60 to 0.25. Therefore, our models can substantially reduce the likelihood of false positives in cases identified as mutated. On top of that, this study compared cell importance in predicting gene mutations using MIL and suggested blasts are crucial indicators for mutation detection. In WSIs with mutations, MIL increased blast cell representation significantly, while in standard WSIs, other cell types increased as blast cells decreased.

In conclusion, our research underscores the immense potential of ensemble learning, upsampling techniques, and MIL in predicting gene mutations in AML patients. Importantly, we accomplished this using training data labeled exclusively at the slide level, eliminating the labor-intensive manual annotation required at the patch and cell levels. This approach streamlined the end-to-end training and prediction process, emphasizing the value of integrating advanced machine-learning approaches to effectively address complex real-world challenges in the field of medical image analysis.

## 5. Acknowledgements

We express our sincere gratitude to the laboratory department staff at NTUH for their exceptional management of the slides. We are grateful to the National Center for High-performance Computing (NCHC) for providing computational and storage resources.

