## Supplementary material for "Annotation-Free Deep Learning for Predicting Gene Mutations from Whole Slide Images of Acute Myeloid Leukemia": Figure S1, S2

### 1. Supplementary Document

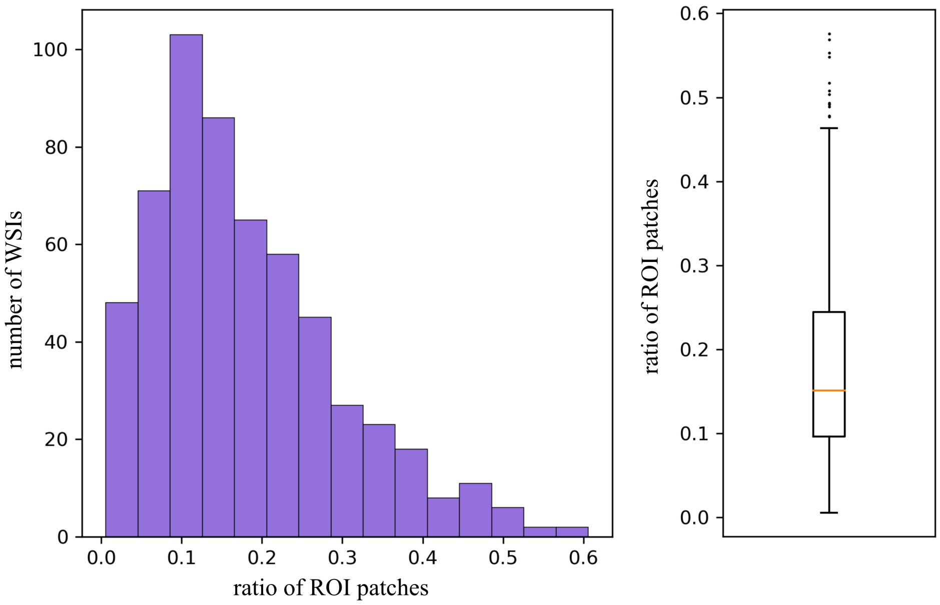


**Figure S1. Distribution (histogram and boxplot) of the selected ROI patches.**

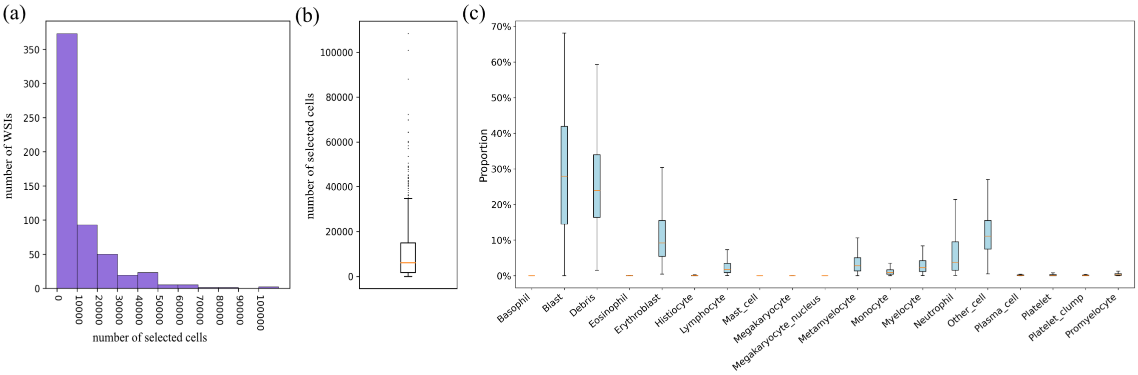


**Figure S2. Distribution of the selected cells. a.** Histogram of the selected cells. **b.** Boxplot of the selected cells. **c.** Proportion for each cell type.
